# A Neurodynamic Model of Inter-Brain Coupling in the Gamma Band

**DOI:** 10.1101/2022.04.08.487686

**Authors:** Moreau Quentin, Adel Lena, Douglas Caitriona, Ranjbaran Ghazaleh, Dumas Guillaume

**Affiliations:** Precision Psychiatry and Social Physiology Laboratory (PPSP), CHU Sainte-Justine Research Center, Montréal, Québec, Canada; Department of Psychiatry, University of Montréal, Québec, Canada; Integrated Program in Neuroscience, McGill University, Montréal, Québec, Canada; Division of Social and Transcultural Psychiatry, McGill University, Montréal, Québec, Canada; Lady Davis Institute for Medical Research, Montreal Jewish General Hospital, Montréal, Québec, Canada; Mila, University of Montréal, Québec, Canada; Human Brain and Behavior Laboratory, Center for Complex Systems and Brain Sciences, Florida Atlantic University, Boca Raton, Florida, USA

**Keywords:** Kuramoto model, Cross-Frequency coupling, hyperscanning, EEG

## Abstract

The use of EEG to simultaneously record multiple brains (i.e., hyperscanning) during social interactions has led to the discovery of inter-brain coupling (IBC). IBC is defined as the neural synchronization between people and is considered to be a marker of social interaction. IBC has previously been observed across different frequency bands, including Theta [4-7 Hz]. Given the proximity of this frequency range with behavioral rhythms, models have been able to combine IBC in Theta with sensorimotor coordination patterns. Interestingly, empirical EEG-hyperscanning results also report the emergence of IBC in the Gamma range [>30 Hz]. Gamma oscillations’ fast and transient nature makes a direct link between Gamma-IBC and other (much slower) interpersonal dynamics difficult, leaving Gamma-IBC without a plausible model. However, at the intra-brain level, Gamma activity is coupled with the dynamics of lower frequencies through cross-frequency coupling (CFC). This paper provides a biophysical explanation for the emergence of Gamma inter-brain coupling using a Kuramoto model of four oscillators divided into two separate (brain) units. By modulating both the degree of inter-brain coupling in the Theta band (i.e., between-units coupling) and CFC (i.e., intra-unit Theta-Gamma coupling), we provide a theoretical explanation of the observed Gamma-IBC phenomenon in the EEG-hyperscanning literature.

## Introduction

Social interaction is a core feature of human life. However, the neural mechanisms that support our capacity to interact with others remain poorly understood due to the fact that neuroscience has mainly focused on recording single participants in isolation rather than assessing several interacting agents simultaneously. Recently, however, the simultaneous recording of multiple brains, commonly known as hyperscanning, has become a popular method within the field of social neuroscience to study interpersonal brain dynamics (Montague et al., 2002; Babiloni et al., 2006; Dumas et al., 2010; Czeszumski et al., 2020). Specifically, electroencephalography (EEG) hyperscanning led to the report of a phenomenon called interbrain coupling (IBC, but see also similar terms such as Inter-brain Synchrony/Synchronization (Dumas et al., 2010; Reinero et al., 2020; Novembre and Iannetti, 2021), a temporal synchronization of neural signals across brains when participants interact (Lindenberger et al., 2009; Dikker et al., 2017; Dikker et al., 2019; Koban et al., 2019). Inter-brain coupling is now widely accepted as a marker of social engagement and successful interpersonal communication, despite the doubt regarding its epiphenomenal nature not being completely lifted (Hamilton, 2020; Holroyd, 2022). IBC has been mostly highlighted using phase synchrony indices such as the Phase-Locking Value (PLV; Lachaux et al., 1999), the Phase-Locking Index (PLI; Tass et al., 1998) and the Partial Directed Coherence (PDC; Baccalá and Sameshima, 2001). This revealed a variety of inter-brain synchronizations across different frequency bands, including in the Theta [4-7 Hz] (Lindenberger et al., 2009; Astolfi et al., 2011; Sänger et al., 2012; Kawasaki et al., 2013) and the Alpha/Mu [8-13 Hz] ranges (Dumas et al., 2010; Naeem et al., 2012; Konvalinka et al., 2014). According to the laws of coordination dynamics, behavioral rhythms of participants during an interaction can both influence and be reciprocally influenced by the behavior of the partner, resulting in a convergence of the dyad’s behavioral rhythms towards a common frequency (Kelso et al., 2013; Tognoli and Kelso, 2015). Given the proximity of Theta and Alpha/Mu frequencies with the rhythms of behavioral sensorimotor coordination, IBC in these ranges can be modeled according to the same coordination dynamics principles and reciprocal exchanges of information across members of an interaction, leading to the brain-behavior coordination dynamics framework (Dumas et al., 2010, 2012a; Kelso et al., 2013; Tognoli et al., 2007; Tognoli & Kelso, 2015).

However, inter-brain synchronizations in higher frequencies such as in the Gamma range [>30 Hz] have also been reported (Dumas et al., 2010; Kinreich et al., 2017; Mu et al., 2017; Barraza et al., 2020). Gamma waves are fast and ultra-fast transient oscillations believed to support local computation (Fries, 2005; Hughes, 2008; Fries, 2009; Magri et al., 2012). Hence, the time scale of this frequency band cannot be directly attributed to behavioral coordination rhythms, leading some to question the validity of observed Gamma-IBC (Holroyd, 2022). On the other hand, at the intra-brain level, an increase of local Gamma amplitude is supported by the phase of lower frequencies (Theta-Gamma coupling) through cross-frequency coupling (CFC; Canolty and Knight, 2010; Lisman and Jensen, 2013; Akkad et al., 2021). CFC has been described as a physiological mechanism capable of coordinating neural dynamics across spatial and temporal scales, where the firing of local neural populations is controlled by larger whole-brain dynamics (Aru et al., 2015). Based on these characteristics, we propose that previously observed Gamma IBC during social interactions can be explained by the combination of two neurophysiological occurrences: 1) inter-brain coupling of lower frequency waves according to coordination dynamics and 2) intra-brain level cross-frequency coupling.

## Results and Discussion

### Schematic Model

To test our hypothesis that IBC in Gamma can be accounted for by the joint effects of Theta IBC and Theta-Gamma CFC, we conceptualized a simple computational model of coupled oscillators. We opted for a model capable of capturing the elementary principles of intra and inter-brain coupling with minimal features. As illustrated in Figure 1A, our model contains two brains, represented as two separate units (units A and B), that are coupled together through inter-brain coupling in the Theta band (θ), while within each unit Theta and Gamma (ɣ) are coupled through CFC.

**Figure 1.**
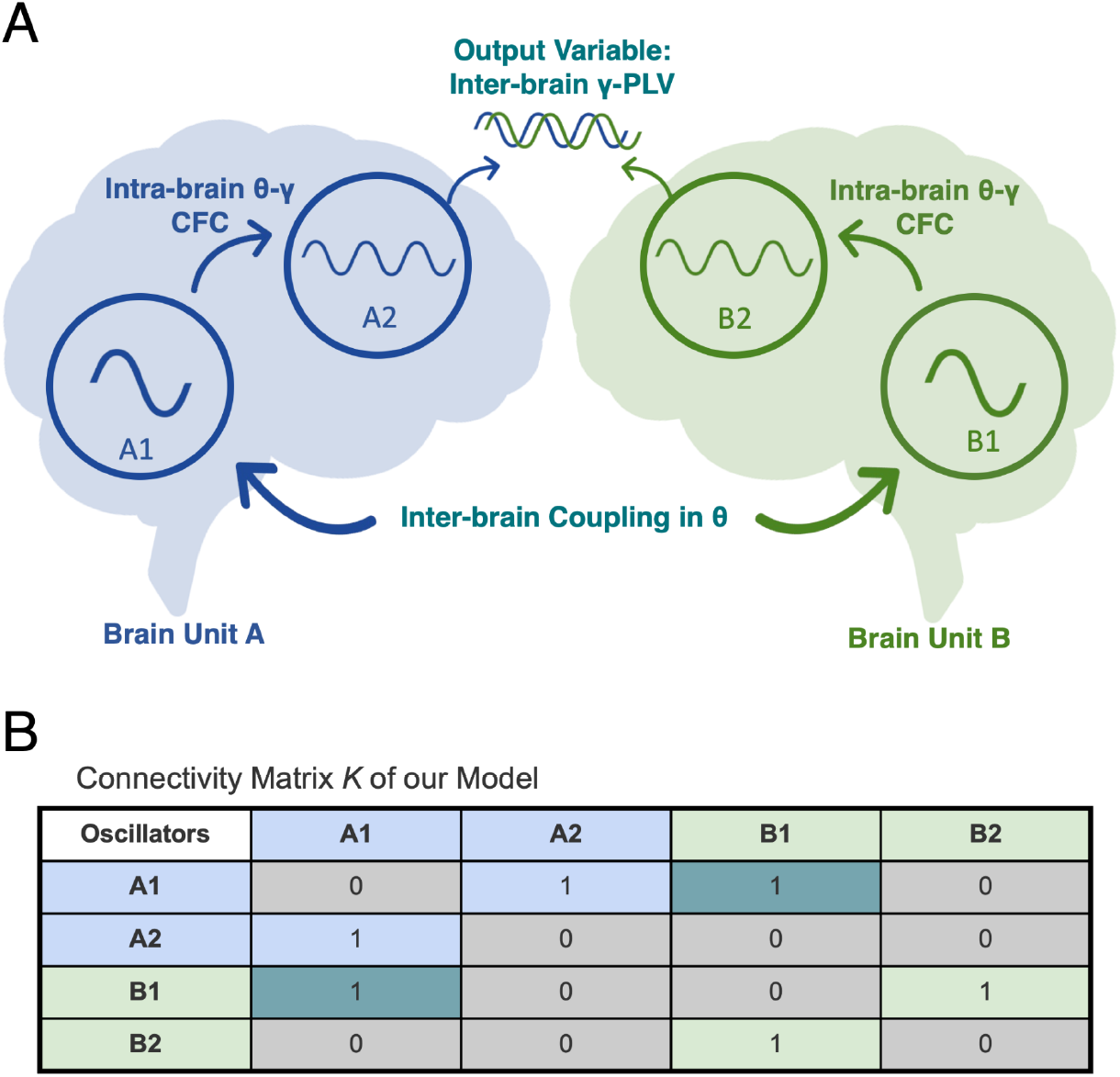
Overview of the model. **A**. Schematic representation of our two-brain model, capable of capturing the elementary principles of intra (A1-A2 and B1-B2) and inter-brain (A1-B1) coupling; **B**. the connectivity matrix K, where 1 means the presence and 0 the absence of a coupling between the oscillators.

### Kuramoto Simulations and Signal-to-Noise ratio

Previous studies used the Kuramoto model for weakly coupled oscillators (Kuramoto, 1975; Acebrón et al., 2005) to demonstrate the effect of intra-brain anatomical and functional connectivity on IBC (Dumas et al., 2012), as well as interpersonal behavioral synchronization strategies and how they rely on the relationship between intra- and inter-unit coupling (Heggli et al., 2019). Generally, the Kuramoto model describes a system of coupled oscillators where the individual oscillators are attracted and entrained to the average rate (in our case, this refers to phase convergence rather than frequency convergence (Cumin & Unsworth, 2007; Dumas et al., 2012). Here, we implemented our model using Kuramoto oscillators, following the connectivity matrix K (Figure 1B). The mean frequency of the oscillators A1 and B1 was set at 6Hz (± 1, i.e., within the Theta range), and the mean frequency of the oscillators A2 and B2 was set at 40Hz (± 1, i.e., within the Gamma range) and without time delays. We simulated time series with a length of 40s (by steps of 10ms). The inter-brain coupling between A1 and B1 in the Theta band and the intra-brain Theta-Gamma CFC (between A1-A2 and B1-B2) were programmed to range from 0 to 1, by steps of 0.1. Simulations were run 10000 times to obtain stable results. We used a Gaussian noise of 0.6 to have a Signal-to-Noise ratio (SNR, see computation in the Methods) of 6.575 dB, which is comparable to SNR found in the EEG literature (Goldenholz et al., 2008).

### Inter-brain Gamma Connectivity

To estimate inter-brain connectivity between the simulated time series of oscillator A2 and B2, we computed the Phase Locking Value (see Material and Methods section). The γ-PLV matrix containing the inter-brain connectivity values between the oscillators A2 and B2 is illustrated by the heatmap in Figure 2.

**Figure 2.**
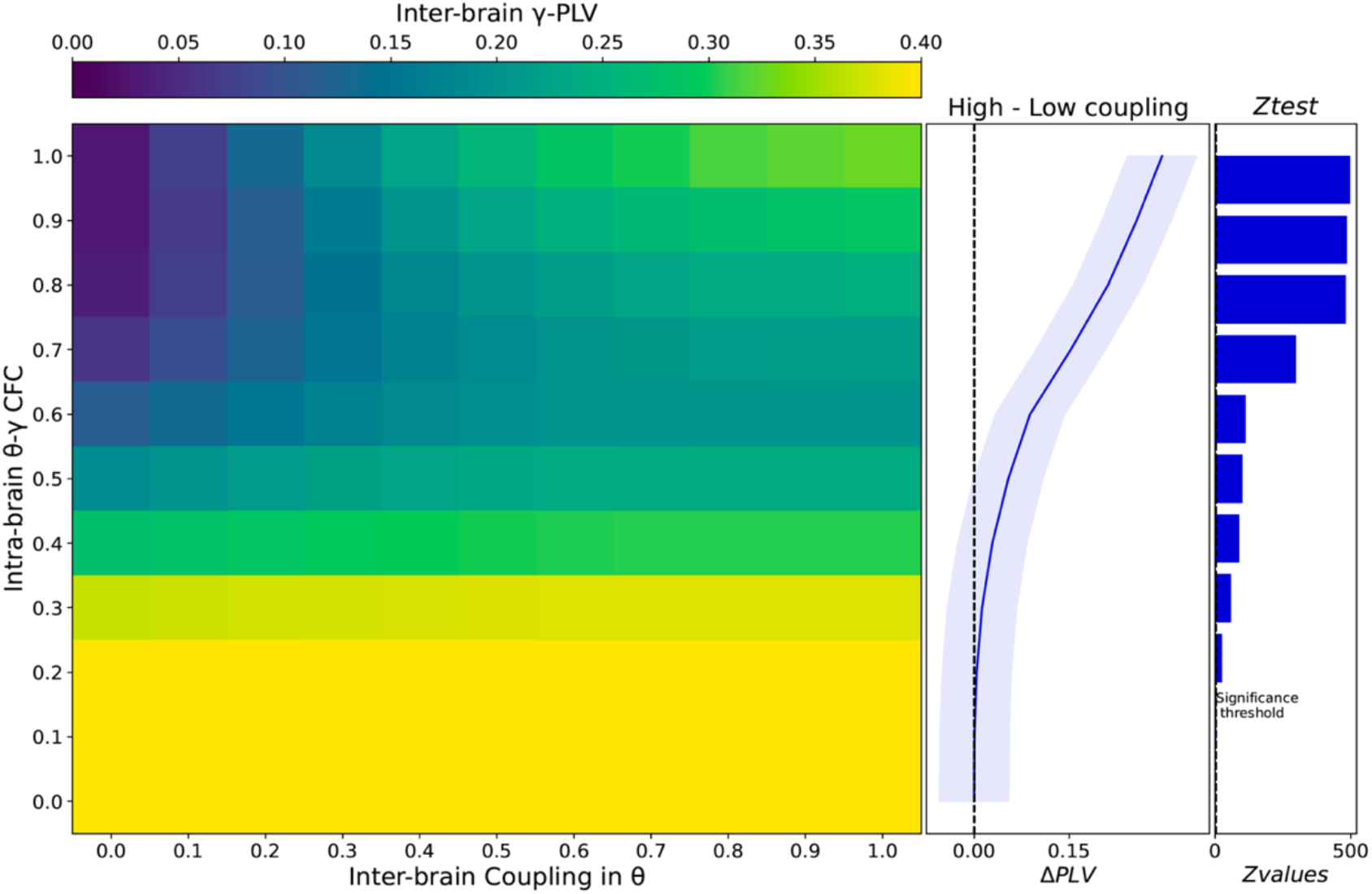
Effect of inter-brain coupling in θ and θ-γ cross-frequency on inter-brain coupling in Gamma. PLV scores between A2 and B2 oscillators reveal that a joint increase of inter-brain coupling in θ and θ-γ cross-frequency coupling account for the observation of inter-brain coupling in the γ band. Panels on the right show that subtracting high θ-IBC and low θ-IBC (i.e., ΔPLV) and Z-testing the values against 0 confirms the pattern observed on the heatmap.

The first observation is that the modulation of θ inter-brain coupling alone has no effect on the γ-PLV values. This is shown by consistent high PLV scores along the x-axis on the heatmap in Figure 2, associated with low values of θ-γ CFC (i.e., between 0 and 0.2). These IBC-independent γ-PLVs represent spurious connectivity that can be explained by the similar properties of the oscillators in our model.

Our second crucial finding is that the increase of θ-γ CFC (above 0.3 on the y-axis of the heatmap in Figure 2) together with an increase of Inter-brain coupling in the θ band is associated with higher PLV scores (see top-right corner values of the heatmap in Figure 2). Additionally, we subtracted the values (i.e., ΔPLV) with the highest degree of IBC (i.e., Interbrain coupling in the Theta band = 1) from the values with the lowest degree of coupling (i.e., Inter-brain coupling in the Theta band = 0) and performed a one-sample z-test (N = 10000) on ΔPLV values against 0 for each values of CFC, confirming incremental effect of CFC on γ-PLV (see bar plots in Figure 2): Z_CFC=0_ = 0.572, *p* = .283; Z_CFC=0.1_ = 4.963, *p* > .001; Z_CFC=0.2_ = 24.421, *p* > .0001; Z_CFC=0.3_ = 58.554, *p* > .0001; Z_CFC=0.4_ = 87.363, *p* > .0001; Z_CFC=0.5_ = 99.601, *p* > .0001; Z_CFC=0.6_ = 112.079, *p* > .0001; Z_CFC=0.7_ = 296.298, *p* > .0001; Z_CFC=0.8_ = 480.977, *p* > .0001; Z_CFC=0.9_ = 484.854, *p* > .0001; Z_CFC=1_ = 495.701, *p* > .0001. These results highlight the impact of the joint increase of θ inter-brain coupling and θ-γ crossfrequency coupling on γ-PLV.

### Biophysical explanation for Gamma inter-brain coupling

Our model provides a biophysical explanation for Gamma inter-brain coupling, an observation that has often been reported in human EEG hyperscanning studies. While IBC in lower frequencies such as Theta and Alpha is in line with brain-behavior coordination dynamics (Dumas et al., 2010, 2012a; Kelso et al., 2013; Tognoli et al., 2007; Tognoli & Kelso, 2015), doubts regarding the validity and the reliability of observed Gamma inter-brain coupling and its potential epiphenomenological nature (i.e., Gamma IBC not actually being a neural correlate of interaction) are not completely lifted (Holroyd, 2022).

Based on evidence from intra-brain, inter-brain, and computational connectivity studies (Dumas et al., 2010; Dumas et al., 2012; Heggli et al., 2019; Loh and Froese, 2021), but also recent account of inter-brain correlations in animals including bats (Zhang and Yartsev, 2019) and mice (Kingsbury et al., 2019; Kingsbury and Hong, 2020), we hypothesized that Gamma IBC could be explained and modeled according to two distinct neurophysiological processes, namely inter-brain coupling in Theta (Lindenberger et al., 2009; Astolfi et al., 2011; Sänger et al., 2012; Kawasaki et al., 2013) and intra-brain Theta-Gamma cross-frequency coupling (Canolty and Knight, 2010; Lisman and Jensen, 2013). First, we showed that our 4-oscillator Kuramoto model, divided into 2 separate units, was able to replicate the core characteristics of both CFC and IBC. Furthermore, our model confirms the hypothesis that IBC in Gamma can be ascribed to intra-brain Theta-Gamma cross-frequency and Theta interbrain coupling, by showing higher PLV scores during the joint increase of both parameters. Hence, our simulations give a biophysical model to the observed Gamma-IBC in the EEG-Hyperscanning literature (Dumas et al., 2010; Kinreich et al., 2017; Mu et al., 2017; Barraza et al., 2020). Our results also illustrate the importance of both intra-brain and inter-brain factors in hyperscanning.

Altogether, through computational modeling, our approach and results advance our mechanistic understanding of IBC, which is crucial to reaching a coherent theoretical framework describing causal relations between socio-cognitive factors, behavioral dynamics, and neural mechanisms involved in multi-brain neuroscience (Moreau and Dumas, 2021).

## Material and Methods

### Dynamical Model of Gamma IBC with Kuramoto

Leveraging Python implementation of Kuramoto systems (Laszuk, 2017), we implemented our model in Python 3.7 (Van Rossum and Drake, 2009) using libraries such as Numpy (Harris et al., 2020), and SciPy (Virtanen et al., 2020) for the computational analyses, and Matplotlib (Hunter, 2007) for the visualization. The Kuramoto model also holds several assumptions: that all oscillators are identical, that the oscillators are innately coupled, and that the oscillations follow a sinusoidal pattern (Kuramoto, 1975; Acebrón et al., 2005; Cumin and Unsworth, 2007; Hudson et al., 2021). Finally, the phase θ of an oscillator i at time t is described by the following dynamical equation:

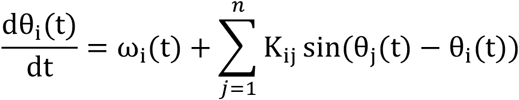

where Kij is the coupling matrix with coupling from oscillator i to oscillator j and ωi is the frequency of oscillator i.

As mentioned above, our model is composed of 4 oscillators, two in each Brain unit (oscillators A1 and A2 in Brain unit A and B1 and B2 in Brain unit B, see Figure 1). The connectivity matrix K is illustrated in Figure 1B. The inter-brain coupling between A1 and B1 in the Theta band and the intra-brain theta-gamma CFC (between A1-A2 and B1-B2) were programmed to range from 0 to 1, by steps of 0.1.

### Inter-brain coupling Measure

To quantify the coupling between the A2 and B2 Gamma oscillators, we used the Phase Locking Value (i.e., PLV) which provides a frequency-specific phase synchrony measure between two signals across time (Lachaux et al., 1999) and is widely used in both intra and inter-brain EEG studies (Tognoli et al., 2007; Dumas et al., 2010; Burgess, 2013; Moreau et al., 2020). We applied a Hilbert transform to extract the instantaneous phase of the signals from oscillators A2 and B2 (see Figure 1A) and computed the ɣ-PLV via the following equation:

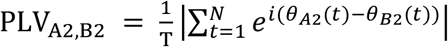

where T is the number of sampled time points and θ_A2_ and θ_B2_ are the instantaneous phase values of oscillators B and D at time point t. PLV values range from 0 to 1, where 0 reflects an absence of phase synchrony and 1 an identical relative phase between the two signals.

### Signal-to-Noise ratio

By extracting the signal and the noise amplitude of the simulated time series, we computed the Signal-to-Noise ratio (SNR) using the following formula:

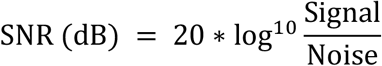

### Data availability

The current manuscript only relies on computational simulations, no data has been recorded. All codes are available at https://github.com/ppsp-team/Hyper-Model. The data file contains the numerical matrices generated to reproduce Figure 2.

